# Ecological predictability emerges at the population level in phytoplankton communities

**DOI:** 10.64898/2026.04.08.717202

**Authors:** Lorenzo Fant, Moritz Klaassen, Onofrio Mazzarisi, Giulia Ghedini

## Abstract

Predicting the composition and dynamics of ecological communities is challenging because complexity increases rapidly with species richness. A common strategy is to adopt a reductionist framework in which community dynamics are inferred from simpler components, such as population-level parameters or organismal traits. However, it remains unclear at which level of biological organization ecological predictability emerges. Here we experimentally test this reductionist cascade in marine phytoplankton communities. We first ask whether multispecies dynamics can be quantitatively predicted from demographic parameters measured in monocultures and species pairs. We then test whether these predictive parameters can themselves be inferred from organismal traits, focusing on cell size. We find that community composition is highly reproducible and can be accurately predicted from population-level parameters measured in simpler experimental settings. In contrast, these parameters do not show systematic relationships with cell size and cannot be predicted from this commonly used trait. These results demonstrate that ecological predictability emerges at the population level, where demographic parameters capture the combined effects of underlying biological processes, but resist further reduction to simple trait-based descriptions, suggesting that ecological interactions reshape organismal performance across levels of organisation.

---

Predicting the diversity and composition of ecological communities is a central challenge in ecology [1, 2]. Yet despite decades of theoretical and empirical work, predicting community composition remains difficult, particularly in species-rich systems where complex interaction networks and stochasticity undermine reproducibility [3, 4].

A major obstacle is the rapid increase in complexity as species richness grows. Each additional species introduces new potential interactions, making multispecies dynamics - and the resulting community composition - increasingly difficult to characterize. In principle, interactions may involve multiple species simultaneously, and the number of such terms increases combinatorially with community size, potentially scaling as *S*^*S*^. Consequently, directly measuring all parameters governing community dynamics becomes infeasible.

Predicting the composition of ecological communities therefore requires some form of reductionism. Rather than attempting to describe all possible interactions, most approaches test the idea that community dynamics can be captured by a limited set of lower-level parameters measured in simpler experimental settings, such as species growth rates and pairwise interaction strengths [5]. Under this framework, community composition and dynamics emerge from the combined effects of these population parameters (Fig. 1).

Empirical tests of this reductionist framework have largely focused on simplified pairwise assembly rules. These approaches rely on pairwise competition experiments to determine which species coexist or exclude one another, and then using these qualitative outcomes to predict coexistence in communities [6, 7]. However, these approaches have produced mixed results, with pairwise outcomes sometimes predicting multispecies coexistence but often failing to do so [7].

The discrepancies between predictions from bottom-up models and observed community coexistence might suggest that ecological dynamics are shaped by higher-order interactions (HOIs) [8], or suffer from strong context dependence [9, 10, 11]. However, recent theory shows that pairwise predictions can fail to predict community assembly even in purely pairwise systems through a mechanism termed *emergent coexistence* [7, 12]. In this case, species that exclude each other in pairwise competition can nevertheless persist collectively due to indirect interaction networks [13], meaning that coexistence cannot always be inferred from qualitative pairwise outcomes alone.

Even if population parameters and pairwise interactions were sufficient, this approach cannot be easily scaled to diverse communities as the number of measurable parameters still increases rapidly with the number of species. One widespread approach is to assume that demographic parameters themselves can be predicted from organismal traits [14, 15, 16]. Under this view, ecological predictability follows a cascade in which community dynamics arise from population processes that may themselves be determined by organismal traits (Fig. 1). Trait-based ecosystem models often exploit this idea by linking physiological parameters to simple traits such as cell size, allowing complex communities to be represented using size-dependent growth and metabolic rates [17, 18, 19].

Body size is a particularly promising candidate because metabolic theory predicts systematic scaling relationships between size, metabolism, and demographic rates [20, 21, 22]. In marine phytoplankton, physiological processes related to energy use —including photosynthesis and respiration— scale allometrically with cell size with an exponent *<* 1 [23]. This means that cell size influences the metabolic expenditure per unit mass of an organism, with potential downstream effects on growth and interactions [16]. However, empirical studies have produced contrasting results regarding the predictive power of body size on ecological performance, including evidence that growth rates may vary weakly with size across phytoplankton taxa [24] and that size differences fail to predict pairwise coexistence [25].

Using phytoplankton communities as a model system, we experimentally test the reductionist cascade linking organismal traits, population parameters, and community dynamics. Rather than relying on qualitative assessments of coexistence (presence/absence), we focus on quantitative predictions of species dynamics over short timescales. Specifically, we test whether the multispecies dynamics during a growth cycle can be predicted from demographic parameters measured in monocultures and species pairs. By predicting the full species dynamics rather than coexistence outcomes, this approach avoids the limitations of qualitative assembly rules.

We then ask whether the demographic and interaction parameters that successfully predict community dynamics can themselves be explained by organismal traits. Focusing on phytoplankton cell size, we test whether size-based scaling relationships predict the parameters governing species growth and competition. This approach provides an integrated test of whether complex ecological dynamics can be predicted from progressively simpler biological descriptions.

**Figure 1.**
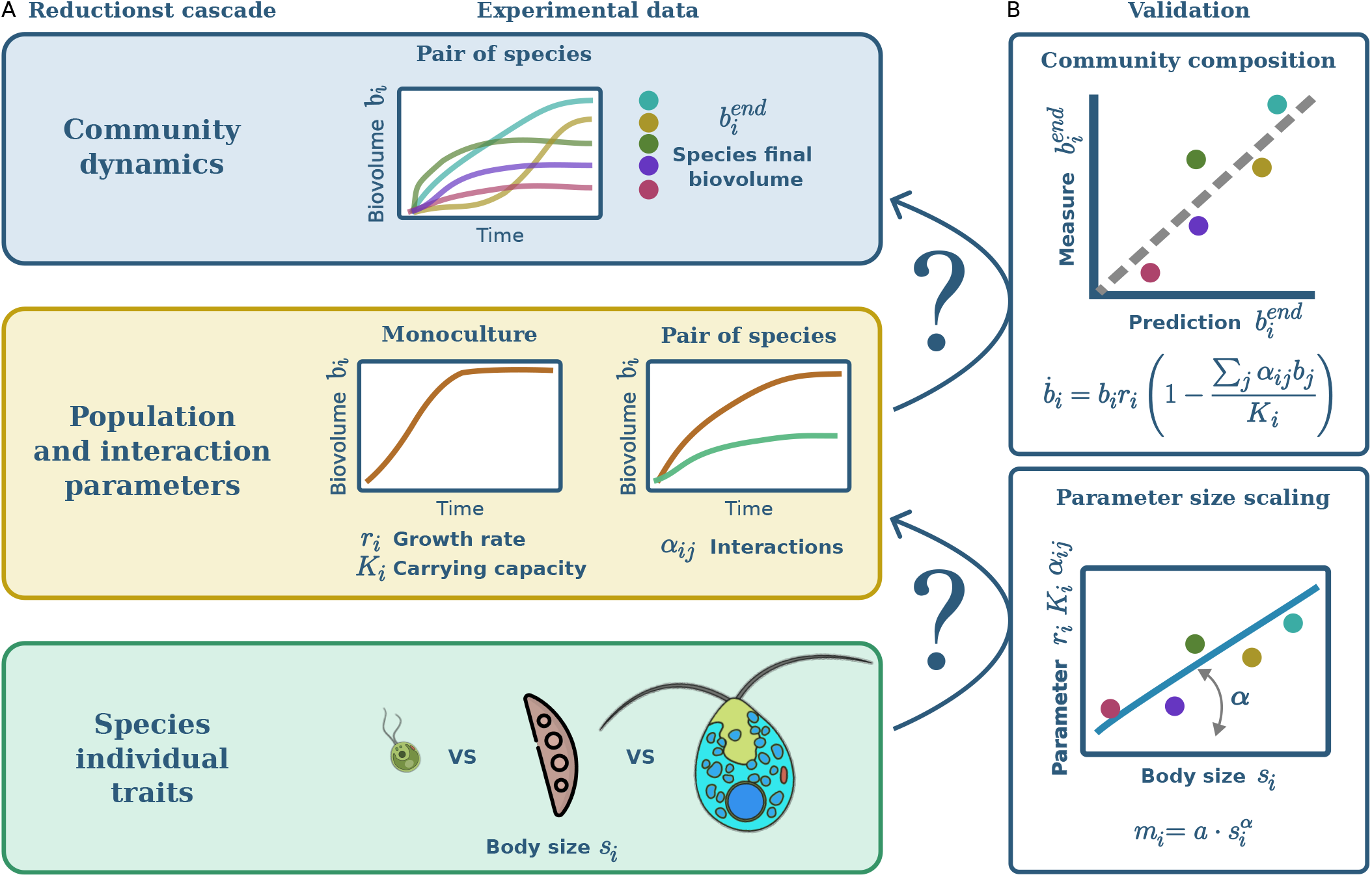
Schematic overview of the predictive cascade tested in this study. We ask whether information measured at one level of biological organisation predicts the next. In panel A we identify the three levels we focus on and the experimental data we collect: 1) individual traits, particularly size *s*_*i*_ given its scaling with energy use → 2) population parameters in monoculture (*r*_*i*_, *K*_*i*_) and interaction coefficients in pairs (*α*_*ij*_) as measures of performance in simple settings → 3) quantitative dynamics and final biovolume composition 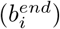 of five-species communities. Panel B shows the analysis we performed to validate the dependence of the level of organisation on the adjacent ones. The validation establish whether the dynamics at one level of organisation can be predicted from the data collected at the lower level.

## Results

### Experimental tests show that community composition is reproducible

Growth trajectories in monocultures, pairs, and five-species communities were highly reproducible across replicates and environmental conditions (Supplementary Figure S1). Monocultures revealed intrinsic differences in productivity among species, reflecting the broad physiological diversity of our species pool. Pairwise experiments demonstrated clear competitive effects, with many species reaching lower final abundances in competition than in isolation, and some inverting the predicted outcomes from monoculture results.

Despite the variability in species abundances and pairwise interactions, five-species communities consistently converged on stable and reproducible compositions within each environmental treatment (Supplementary Figure S1). While light, salinity, and experiment setting modulated both growth rates and competitive outcomes, the rank order of species success was robust across replicates. These results highlight both the determinism, seen in the reproducibility across the replicates, and the complexity, given by the large variance of final species abundances, of phytoplankton community assembly. They provide the empirical foundation for testing whether monoculture and pairwise parameters are sufficient to predict multi-species outcomes.

### Quantitative predictions of community composition

Our goal is to make quantitative, predictions of community dynamics and final composition from measurements performed in simpler settings. Therefore, we infer species demographic traits from monocultures, and pairwise interaction coefficients from two-species cultures, and then use only these inputs to predict the outcomes of five-species assemblages, in the form of final species biovolumes. Crucially, these predictions are strictly out-of-sample: they are evaluated on independent multispecies communities using parameters inferred exclusively from monoculture and pairwise experiments not used in prediction.

We model the dynamcis of the species total biovolume *b*_*i*_ using a Generalized Lotka–Volterra (GLV) form [26, 27]:

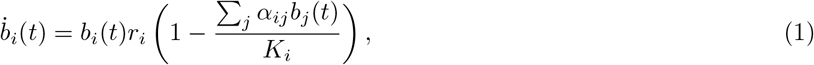

where *r*_*i*_ is the growth rate, *K*_*i*_ is the single-species carrying capacity (in biovolume units), and *α*_*ij*_ quantifies the per-biomass effect of species *j* on species *i* (with *α*_*ii*_ = 1). Crucially, parameter estimation uses only monocultures and pairs, and predictions are evaluated on independent five-species communities.

We do not assume a specific functional form; rather we use GLV because it provides a minimal, interpretable baseline that maps directly onto our measurements and yields falsifiable quantitative predictions. In this context, higher-order interactions are operationally defined as systematic, reproducible deviations between multi-species dynamics and the dynamics predicted from monoculture traits and pairwise coefficients.

#### a. Neutral interactions

As a baseline—and to isolate the predictive information contained in monoculture measurements—we first asked whether species demography in isolation is sufficient to predict competitive outcomes when interactions are assumed to be “neutral” (i.e., independent of competitor identity). Under this assumption, species differ in their intrinsic demographic parameters (*r*_*i*_ and *K*_*i*_), but the *per-biomass* competitive effect of any neighbor is the same.

For each species *i*, we estimated two parameters from monoculture growth: the intrinsic growth rate *r*_*i*_ (low-density exponential phase) and the carrying capacity *K*_*i*_ (the maximum supported biovolume in a given environment). We parameterized these using a logistic model, identical to eq. 1 with one species only.

To predict dynamics in competition, we replaced the population biovolume *b*_*i*_ with total community (or pair) biovolume Σ _*j*_ *b*_*j*_, equivalent to setting *α*_*ij*_ = 1 for all *i, j* in eq. 1 (including *α*_*ii*_ = 1).

Despite ignoring species-specific interaction coefficients, this neutral baseline predicted final biovolumes in pair and multi-species cultures with substantial accuracy (pairs Spearman *ρ* = 0.64, *r*^2^ = 0.61 (0.52-0.68), *p <* 10^−22^, communities Spearman *ρ* = 0.83, *r*^2^ = 0.88 (0.83-0.91), *p <* 10^−41^; Fig. 2A). Thus, in agreement with previous work [28], a large fraction of predictable variation in community composition is already captured by species-level demographic parameters measured in isolation. At the same time, these results show that the data are inconsistent with strict ecological neutrality, in which species are identical: differences in *r*_*i*_ and *K*_*i*_ are necessary to explain the observed heterogeneity in abundances. Discrepancies, particularly evident for *Tisochrysis* in the PT data (Fig. 2A), point to the importance of interactions in this system. In fact, the neutral model consistently predicts a higher final biomass for this species than that observed, both in pairs and in communities, suggesting growth suppression when it grows with other species compared to monoculture conditions. Similarly, the model underestimates the abundance of the dominant species (*Phaeodactylum*).

**Figure 2.**
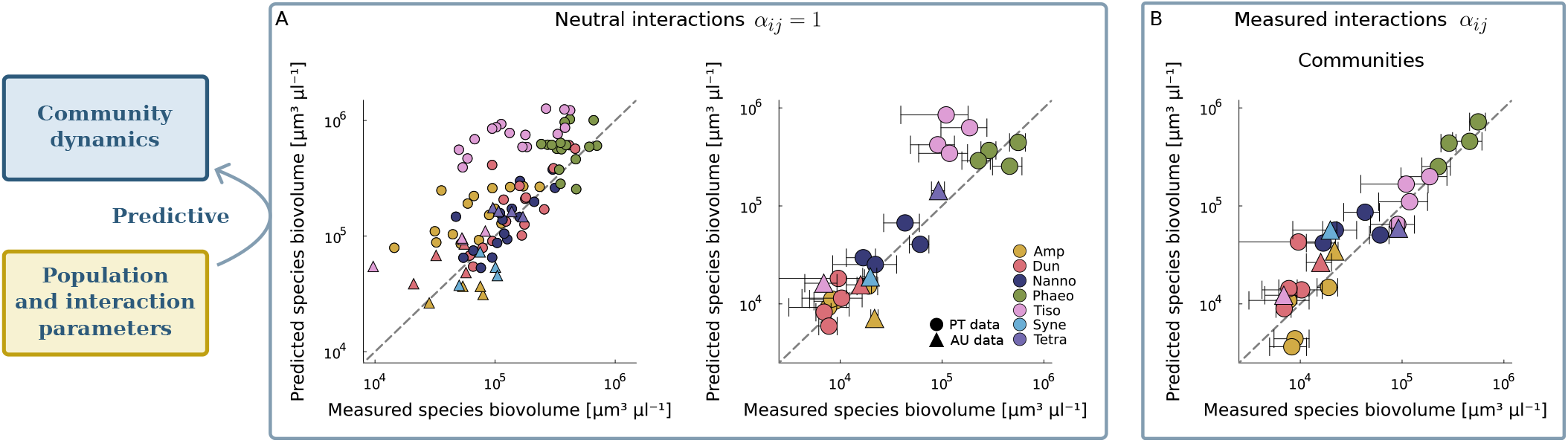
Ecological parameters infered from monocultures and pairs data quantitatively predict community composition. Points compare observed and predicted final biovolume density of each species in the community; the diagonal line indicates perfect prediction. **A** Neutral baseline: monoculture parameters (*r*_*i*_, *K*_*i*_) partially predict pair (left) and community (right) outcomes assuming competitor identity is irrelevant (*α*_*ij*_ = 1). **B** Pairwise model: monoculture parameters together with interaction coefficients inferred from pair cultures (*α*_*ij*_) predict endpoint composition in independent five-species communities.

#### b. Pairwise interactions

A model based on neutral interactions provides a stringent baseline, but it implicitly assumes that all competitors exert the same per-biomass effect. Because phytoplankton species differ in resource acquisition, morphology, and physiology [19], competitive impacts are expected to be species-specific. We therefore use pair cultures to infer interaction coefficients and ask whether these *pairwise* measurements improve quantitative predictions, and— crucially—whether they generalize from two-species cultures to multi-species communities.

We observe that pairs consistently reach lower total biovolume than both monocultures and communities, independent of species composition (Supplementary Figure S2). This pattern has been reported previously and a potential explanation has been proposed: in multispecies communities, species might partition their impacts on others, reducing the effects of strong competitive interactions [29]. Because it is not central to our scope, we do not investigate it further, but account for it explicitly to avoid biasing the inferred interaction coefficients. We therefore estimated a reduction factor for total carrying capacity in pairs by calculating the average ratio between the final total biomass in pairs 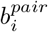 and monocultures 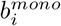 as 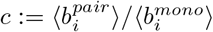. We used an identical factor for all species and treatments, but differing between experimental locations, as these showed to behave quite differently. We then add it to eq. 1 by multiplying it to all the *K*_*i*_, before introducing the competition coefficients *α*_*ij*_.

With *r*_*i*_ and *K*_*i*_ fixed from monocultures and *c* estimated, we parameterized species-specific interaction coefficients *α*_*ij*_ from the two-species time series and used them to integrate the GLV model (Eq. 1) forward in time from the initial conditions of each experimental community. We then compared predicted endpoint biovolumes with observations in independent five-species assemblies.

While monoculture demography alone already captured most of the variance in community biomass (r^2^ = 0.88), it frequently misordered species in the competitive hierarchy (Fig. 2A). Incorporating pairwise interaction coefficients substantially improved the recovery of species rankings, increasing Spearman rank correlation between predicted and observed biovolumes from *ρ* = 0.83 to *ρ* = 0.91 (*r*^2^ = 0.89 (0.85-0.92), *p <* 10^−43^) (Fig. 2B). Improvement was consistent across environments, indicating that interaction identity contains additional, transferable information about community dynamics. Predictive accuracy remained robust across two orders of magnitude in final species biovolume, suggesting that the predictability is not restricted to dominant taxa. Moreover, we see that considering pairwise interactions better identifies the effects of environmental conditions on species growth, in particular for highly abundant species (Fig. 2B and Fig. S3).

Given the wide variability in species abundances within communities, we asked whether community dynamics are effectively driven by only one or a few dominant species, or whether multiple species interactions contribute to the observed outcomes. This distinction is important because if only one or two species dominated the dynamics, the system would not contain enough interacting components to meaningfully assess higher-order effects. To address this, we compared the residual prediction errors (i.e., the distances from the 1:1 dashed line in Fig. 2) with the errors obtained from a simpler baseline in which community outcomes are predicted directly from pairwise experiments, using only single-interaction information [30]. We find (Fig. S4) that predictions based on the full multi-species dynamics are substantially more accurate than those based on single pairwise interactions. This result indicates that multiple species contribute simultaneously to the dynamics, and that community outcomes cannot be reduced to the effect of one or two dominant competitors.

Residual prediction errors provide an empirical window into effects beyond our pairwise description. However, given their extremely low amplitudes, a parsimonious interpretation is that residuals reflect a combination of measurement noise, replicate-to-replicate variability in initial conditions, and the assumed functional form of the GLV model, rather than systematic higher-order effects.

### Size Dependence of Ecological Parameters

The ability of monoculture demographic traits and pairwise coefficients to quantitatively predict multi-species outcomes shows that the inferred parameters capture reproducible, transferable features of the ecological dynamics rather than overfitting idiosyncrasies of a particular dataset. Having established this predictive accuracy, we next asked whether these empirically inferred parameters can themselves be anticipated from a simple organismal trait: body size. We focus on size because metabolic theory predicts systematic scaling of physiological rates with body mass [21, 22], including in this system[23, 31], and metabolism relates to population growth and demography [32, 33, 34]. If size were a dominant driver of competitive ability, then variation in *r*_*i*_, *K*_*i*_, and interaction coefficients should follow monotonic size trends.

We do not find evidence for monotonic correlation between growth rates and cell size (Fig. 3A, *F*_1,23_ = 1.08, *p* = 0.31). Similarly, carrying capacities do not vary systematically with cell size across our dataset (Fig. 3B, *F*_1,23_ = 0.3, *p* = 0.6). At most, intermediate-sized species (∼ 100*µm*^3^) tended to exhibit higher growth rates and carrying capacities, consistent with previous observations [35] on C-fixation and cell abundance [24, 23] and suggestive of trade-offs between the metabolic power of small cells and the efficiency of larger ones [36].

**Figure 3.**
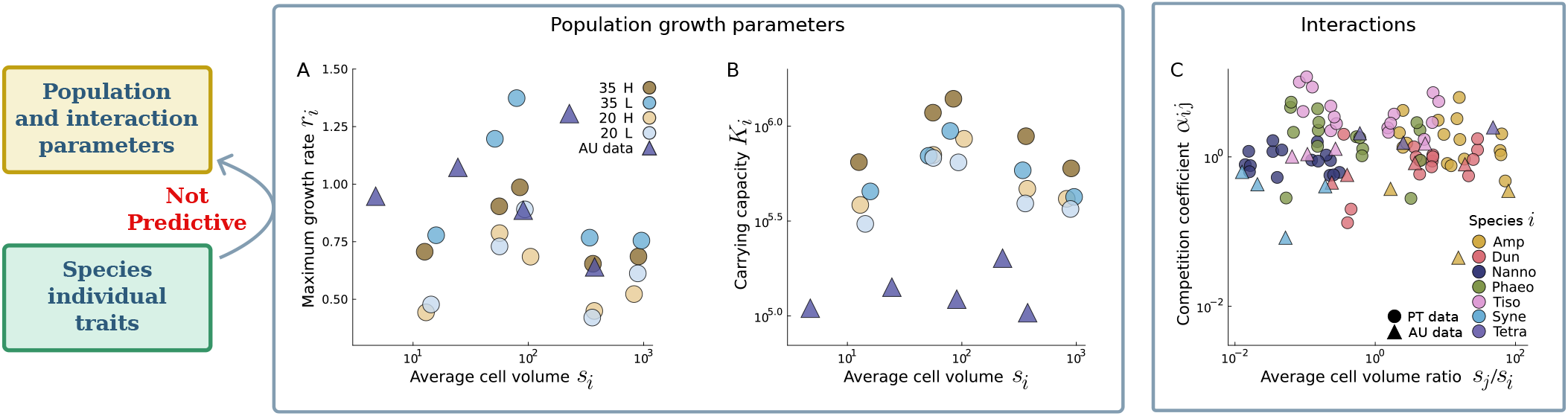
Ecological parameters estimated on species dynamics show no monotonic scaling with cell size. (A,B) Intrinsic growth rates and carrying capacities estimated from monocultures do not correlated with species size. (C) Similarly, pairwise interaction coefficients *α*_*ij*_ show no relationship with the competitors’ size ratio *s*_*j*_*/s*_*i*_.

We next asked whether competitive impacts depend on size asymmetries between interacting species. To avoid bias introduced by absolute size differences, we tested for a relationship between interaction coefficients and the size ratio between competitors. The competition coefficients *α*_*ij*_ showed no detectable dependence on the ratio *s*_*j*_*/s*_*i*_ where *s* is cell size (volume) (Fig. 3C; *F*_1,20_ = 0, *p* = 0.997).

## Discussion

Predicting the composition of ecological communities is challenging because the number of potential interactions grows rapidly with species richness. A common response to this complexity is to adopt reductionist approaches that attempt to describe community dynamics using simpler components. Here we experimentally tested this reductionist cascade by asking whether community dynamics can be predicted from population-level parameters and whether these parameters themselves can be inferred from organismal traits.

Our results show that community dynamics are highly predictable from population-level information. Across environments and experimental replicates, five-species phytoplankton communities converged on reproducible compositions, and their final biovolumes could be accurately predicted using demographic parameters measured in monocultures together with interaction coefficients estimated in species pairs. Because these predictions were evaluated on independent multispecies communities, this success indicates that the inferred parameters capture transferable features of the ecological dynamics rather than idiosyncrasies of particular cultures. More broadly, the reproducibility of trajectories and the quantitative agreement between predicted and observed abundances demonstrate that a minimal model parameterized from simpler experimental settings can provide a robust description of community dynamics.

Although population parameters proved highly informative, we found little evidence that they can be reduced further to simple organismal traits. Metabolic theory predicts systematic scaling relationships between body size, metabolism, and demographic rates [21, 22], leading to the expectation that competitive outcomes in phytoplankton communities might be organized along a size hierarchy. However, neither intrinsic growth rates, carrying capacities, nor interaction coefficients showed monotonic relationships with cell size in our experiments. These results indicate that while population parameters integrate the ecological mechanisms governing species performance, they are not straightforwardly inferable from body size alone.

This discrepancy highlights an important gap in our understanding of the link between organismal traits and ecological dynamics. As species move from isolated conditions to pairwise and multispecies contexts, growth and competition may integrate indirect effects and context-dependent processes that are not apparent at the level of individual traits. One possiblity is that competitive outcomes emerge from multidimensional trait differences rather than a single dominant trait [37]. Phytoplankton species differ not only in size but also in nutrient acquisition strategies, storage capacity, metabolic efficiency, and physiological plasticity [19]. In the presence of interactions, more traits could become important to determine organismal growth and competitive success. This would justify why size, that proved to be highly predictive of individual metabolism and growth[22], cannot capture the performance of species in more realistic ecological scenarios.

Understanding how these mechanisms combine to shape demographic performance therefore remains an important challenge. Future work should investigate which combinations of traits—or which interaction-mediated processes—most strongly determine the population-level parameters that ultimately govern community dynamics. Identifying these links could clarify why simple trait-based approaches often fail and how organismal biology translates into ecological interactions.

Our conclusions should also be interpreted in the context of the experimental system. Phytoplankton communities are primarily structured by competition for essential resources such as light and nutrients, which may limit the scope for complex interaction structures compared with systems involving metabolic cross-feeding or strong trophic feedback [38]. In addition, although five-species communities already represent substantial complexity for controlled laboratory experiments, future work should examine how predictive accuracy scales with increasing diversity, where weak indirect effects might accumulate at higher richness and eventually erode predictability.

Together, our results reveal a clear asymmetry in the reductionist cascade linking traits, populations, and communities. Population-level and interaction parameters inferred from simple species dynamics provide a robust and predictive description of community composition, whereas simpler organismal traits such as body size do not reliably capture the ecological differences that determine competitive outcomes. This suggests that demographic parameters and interactions occupy an intermediate but particularly informative level of description: they integrate multiple physiological and ecological processes while remaining sufficiently simple to support quantitative prediction of multispecies dynamics.

More broadly, our findings illustrate that ecological communities can be more predictable than their apparent complexity suggests, provided that the appropriate level of description is used. Rather than attempting to predict community dynamics directly from organismal traits or from exhaustive multispecies interaction networks, our results suggest that population-level demographic parameters—measurable from monocultures and pairs—provide a sufficient and experimentally tractable basis for quantitative community prediction.

## 1 Materials and Methods

### Experiments Overview

To test these ideas, we use data collected in two separate experiments[39, 28]. Briefly, in each experiment we sourced five phytoplankton strains from local culture collections (ANACC in Australia, Roscoff in France) and maintained these species in the laboratory for several months at constant light and temperature conditions before starting experiments. The 10 strains we used in total span major phytoplankton functional groups and nearly three orders of magnitude in cell size, thereby capturing broad ecological and physiological diversity (Supplementary Table S1).

Within each experiment, we quantified growth under three experimental configurations: monocultures, species pairs, and five-species communities. We tracked the cell size and abundance of species in each culture from low density until carrying capacity, which we used to calculate species biovolume density - a proxy for biomass density[40]. Experiments were performed across five distinct environmental conditions differing in light intensity, salinity, and bubbling regime. Monocultures provided baseline parameters for intrinsic growth, while pairwise cultures revealed competitive interactions between species. We then assess whether these parameters predict the final composition of five-species communities across environmental conditions.

By focusing on reproducibility and quantitative prediction of species abundances, we directly evaluate whether pairwise interactions are sufficient to explain community outcomes [6, 41]. The five-species communities served as the critical test of our predictive framework, allowing for the potential indirect effects that could emerge in multispecies communities. Each configuration (monoculture, pairs, communities) was replicated across all environmental conditions, enabling us to assess both the reproducibility of outcomes and the robustness of species interactions under contrasting resource availability and stress. Finally, we test if the parameters that describe species performance and interactions scale predictably with body size. If this relationship holds, predictions of community composition could be drawn directly from simpler organismal traits without the need to measure interactions within and between species.

### Phytoplankton culturing and sampling

We sourced five marine phytoplankton strains from the Australian National Algae Culture Collection in Australia and five from the Roscoff Culture Collection in France. Experiments with these strains were conducted at Monash University (Australia) and the Gulbenkian Institute (Portugal), respectively. Within each location (hereafter termed AU and PT), species were grown as monocultures, pairwise combinations, and communities containing the five local species together, in temperature-controlled rooms at 22 ± 1^◦^C using standard f/2 enriched seawater medium [42]. Changes in the biovolume of each species were tracked over time for 10 and 16 days in the two locations, respectively.

### Australia experimental setup

The strains obtained from the Australian National Algae Culture Collection were: *Amphidinium carterae* (CS-740), *Tetraselmis* sp. (CS-91), *Dunaliella tertiolecta* (CS-14), *Tisochrysis lutea* (CS-177), and *Synechococcus* sp. (CS-94). We used these strains to establish monocultures (*n* = 3 per strain), pairwise combinations (*n* = 3 per pair), and five-species communities (*n* = 5), for a total of 50 cultures. Each strain was cultured individually in 2 L glass bottles for two months prior to the experiment.

During the experiment, which lasted 10 days, cultures were grown in 200 ml clear culture flasks filled to 100 ml under a 14:10 h light–dark cycle at non-saturating irradiance levels (115 ± 5 *µ*mol m^−2^ s^−1^). Flasks were shaken and randomly rearranged on shelves daily. Nutrients were added daily by replacing 10% of the medium from each flask (used for sampling) with fresh f/2 medium, corresponding to a dilution rate of 0.1 day^−1^.

We adopted a substitutive design, such that all cultures were inoculated with the same total biovolume of approximately 6 × 10^8^ *µ*m^3^ (approximately 10^3^ *µ*m^3^ *µ*l^−1^), with equal initial biovolume contributions from each species within treatments. Changes in species biovolume were determined using light microscopy (see Sampling of species biovolume below). Cultures were sampled daily for the first five days and every second day thereafter for a total of eight sampling times (days 0, 1, 2, 3, 4, 6, 8, and 10). Further details are provided in [39].

### Portugal experimental setup

The strains obtained from the Roscoff Culture Collection were: *Amphidinium carterae* (RCC88), *Dunaliella tertiolecta* (RCC6), *Phaeodactylum tricornutum* (RCC2967), *Tisochrysis lutea* (RCC90), and *Nannochloropsis granulata* (RCC438). These species were grown either alone in monoculture, in pairwise combinations, or together in a five-species community for 16 days under two levels of salinity (ambient 35 ppt or reduced 20 ppt) and light intensity (60 or 30 *µ*mol photons m^−2^ s^−1^) in cross combination to simulate a gradient of stressful environments.

For each salinity–light combination we established five replicate communities (equal initial biovolumes of the five species; *N* = 20 total), two replicate monocultures of each species (*N* = 40), and two replicates of each species pair (*N* = 80) in glass bottles filled to 200 ml. Pair cultures were grown and sampled at a separate time for logistical reasons and were not included in [28].

The position of cultures was randomized at each sampling day and cultures were continuously bubbled for mixing. Cultures were maintained under a 12:12 h light–dark cycle. Initial total biovolume was approximately 4 × 10^9^ *µ*m^3^ (approximately 10^3^ *µ*m^3^ *µ*l^−1^) for each treatment. Changes in species abundance, cell size, and biovolume were tracked through microscopy as described below. Unlike the AU experiment, medium was not replenished during the PT experiment.

Monocultures and communities were sampled 6 and 7 times, respectively, over the 16-day experiment (days 2, 5, 7, 9, 13, and 15 for monocultures; days 2, 3, 7, 9, 12, 14, and 16 for communities). Pair cultures were sampled on alternate days.

### Sampling of species biovolume

In both experiments, 1 ml samples from each culture were fixed with 1% Lugol’s solution to quantify cell size and density. From each fixed sample, 10 *µ*l were loaded onto a Neubauer improved counting chamber and photographed using an Olympus IX73 inverted microscope at 400× magnification.

Images were processed in Fiji/ImageJ to quantify cell volume (*µ*m^3^), cell density (cells *µ*l^−1^), and species biovolume, calculated as the product of these two quantities (*µ*m^3^ *µ*l^−1^). Cell volume was estimated from the major and minor axes of each cell by assigning species-specific geometric shapes [43]. Most species were approximated as prolate spheroids, whereas *Synechococcus, Tisochrysis*, and *Nannochloropsis* were approximated as spheres. Total biovolume in species mixtures (pairs or communities) was calculated as the sum of the biovolumes of individual species.

### GLV parameterization

We parameterized a generalized Lotka–Volterra (GLV) model in two stages using monoculture and pairwise time series, and then integrated the resulting multispecies dynamics to predict final community composition.

For each species (and for each PT treatment), intrinsic growth parameters were estimated from monoculture biovolume trajectories by fitting a logistic curve:

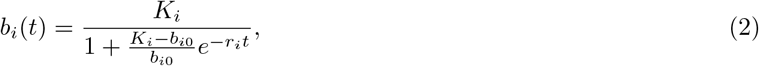

where *b*_*i*_ is species biovolume, *µ*_*i*_ is maximum growth rate, and *K*_*i*_ is carrying capacity.

Pairwise competition coefficients were inferred by jointly fitting both species in each pair trajectory, while keeping monoculture-derived parameters fixed. The fitted discrete-time update for species *i* in pair (*i, j*) was:

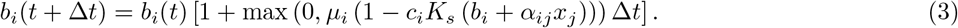

Here, *α*_*ij*_ is the effect of species *j* on species *i*, and *c* is a dataset-level scaling factor estimated from the ratio between final pair and final monoculture biovolume.

For each pair, (*α*_*ij*_, *α*_*ji*_) were estimated by nonlinear least squares over all replicate trajectories for that pair. This produced full interaction matrices with diagonal terms fixed to neutral self-interaction (*α*_*ii*_ = 1): PT as a treatment-specific tensor (4 × 5 × 5), and AU as a single 5 × 5 matrix.

### Community prediction by GLV integration

Using the estimated interaction matrices and monoculture parameters, we predicted final 5-species community composition by forward integration of:

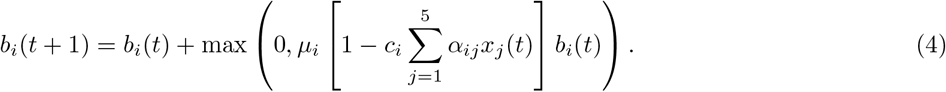

The integration is performed in discrete time as data are collected at most once a day. For PT, *µ*_*i*_ and *c*_*i*_ were treatment-specific means across monoculture replicates. For AU, species-level means were used. Initial conditions were set from the earliest measured community biovolumes, and integration proceeded to the last observed day in each experiment.

To isolate the role of inferred interactions, we also ran a neutral baseline with *α*_*ij*_ = 1 for all (*i, j*). Predicted final biovolumes from both models were compared against measured final community composition (log-scale correlations and scatter comparisons).

### Rescaling of carrying capacities when fitting pairwise interactions

Pairwise interaction coefficients were estimated using a generalized Lotka–Volterra (GLV) framework parameterized from monoculture demographic measurements. In this model, the carrying capacity *K*_*i*_ measured in monoculture represents the equilibrium biomass species *i* would reach in isolation under the experimental conditions. However, when two species were grown together, the total biomass reached at the end of the growth cycle was systematically lower than the biomass obtained in monocultures.

This pattern is illustrated in Fig. S2. In both experimental locations, cultures containing two species reached lower total biovolumes than monocultures grown under the same conditions. In contrast, multispecies communities reached total biovolumes comparable to monocultures. Thus, pair cultures appear to experience a systematic reduction in realized total biomass relative to the monoculture carrying capacities.

To account for this effect when estimating interaction coefficients, we introduced a simple rescaling factor applied to the carrying capacities when fitting the pairwise data. Specifically, we defined a reduction factor

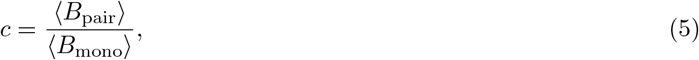

where ⟨*B*_pair_⟩ is the mean final total biovolume across all pair cultures and ⟨*B*_mono_⟩ is the mean monoculture carrying capacity across species. When fitting pairwise interactions, monoculture carrying capacities were multiplied by this factor, effectively adjusting the expected total biomass in two-species systems while preserving the relative demographic differences among species.

This correction prevents systematic overestimation of equilibrium biomass when applying monoculture parameters to pair cultures and allows interaction coefficients to capture deviations in relative species performance rather than differences in overall biomass production. Importantly, the rescaling affects only the global biomass scale and does not alter the inferred competitive hierarchy among species.

Because multispecies communities did not show the same systematic biomass reduction, the rescaled carrying capacities were used only when fitting pairwise interaction coefficients, while predictions for multispecies communities used the original monoculture-derived demographic parameters.

Finally, this rescaling does not introduce additional degrees of freedom in the inference of interaction coefficients. The factor *c* is computed directly from the observed mean biomass in pair and monoculture treatments and is not fitted to improve model agreement with pairwise outcomes. It therefore corrects a systematic shift in total biomass production between experimental treatments without altering the structure of inferred species interactions.

### Testing correlations with cell size

We used a simple linear model in R (version 4.4.2) to test if cell size explains the variability in growth rates and carrying capacities of our species across all environmental conditions. We did the same analysis for competition coefficients vs size differences. All variable (except growth rates) were log10-transformed to reduce the skewness of the data - this transformation did not change the result (all relationships were not significant even when tested on normal data)

## Supporting information

supplementary materia

## References

[1] J. H. Lawton, “Are there general laws in ecology?,” Oikos, vol. 84, no. 2, pp. 177–192, 1999.

[2] M. Vellend, “Conceptual synthesis in community ecology,” The Quarterly Review of Biology, vol. 85, no. 2, pp. 183–206, 2010.

[3] C.A. Serván and S. Allesina, “Tractable models of ecological assembly,” Ecology Letters, vol. 24, no. 5, pp. 1029– 1037, 2021.

[4] G. E. Hutchinson, “The paradox of the plankton,” The American Naturalist, vol. 95, no. 882, pp. 137–145, 1961.

[5] L. Buche, L. G. Shoemaker, L. M. Hallett, I. Bartomeus, P. Vesk, C. Weiss-Lehman, M. Mayfield, and O. Godoy, “A continuum from positive to negative interactions drives plant species’ performance in a diverse community,” Ecology Letters, vol. 28, no. 1, p. e70059, 2025. e70059 ELE-00947-2024.R1.

[6] J. Friedman, L. M. Higgins, and J. Gore, “Community structure follows simple assembly rules in microbial microcosms,” Nature Ecology & Evolution, vol. 1, no. 5, pp. 1–7, 2017.

[7] C.-Y. Chang, D. Bajić, J. C. C. Vila, S. Estrela, and A. Sanchez, “Emergent coexistence in multispecies microbial communities,” Science, vol. 381, no. 6655, pp. 343–348, 2023.

[8] M. M. Mayfield and D. B. Stouffer, “Higher-order interactions capture unexplained complexity in diverse communities,” Nature ecology & evolution, vol. 1, no. 3, p. 0062, 2017.

[9] I. Billick and T. J. Case, “Higher order interactions in a natural community: I. characterization and importance in the desert winter annual plant community,” Ecology, vol. 75, no. 8, pp. 2291–2302, 1994.

[10] J. M. Levine, J. Bascompte, P. B. Adler, and S. Allesina, “Beyond pairwise mechanisms of species coexistence in complex communities,” Nature, vol. 546, no. 7656, pp. 56–64, 2017.

[11] A. Sanchez-Gorostiaga, D. Bajić, M. L. Osborne, J. F. Poyatos, and A. Sanchez, “High-order interactions distort the functional landscape of microbial consortia,” PLoS biology, vol. 17, no. 12, p. e3000550, 2019.

[12] G. Aguadé-Gorgorió and S. Kéfi, “Emergent coexistence and the limits of reductionism in ecological communities,” PLOS Computational Biology, vol. 22, no. 3, p. e1014116, 2026.

[13] M. Barbier, C. De Mazancourt, M. Loreau, and G. Bunin, “Fingerprints of high-dimensional coexistence in complex ecosystems,” Physical Review X, vol. 11, no. 1, p. 011009, 2021.

[14] L. Chalmandrier, F. Hartig, D. C. Laughlin, H. Lischke, M. Pichler, D. B. Stouffer, and L. Pellissier, “Linking functional traits and demography to model species-rich communities,” Nature Communications, vol. 12, no. 1, p. 2724, 2021.

[15] J. Figueiredo, A. H. Baird, S. Harii, and S. R. Connolly, “Predicting demography from species traits: larval development time and its sensitivity to warming depend on egg size in corals,” Coral Reefs, vol. 45, no. 1, pp. 81–91, 2026.

[16] D. J. Wieczynski, P. Singla, A. Doan, A. Singleton, Z.-Y. Han, S. Votzke, A. Yammine, and J. P. Gibert, “Linking species traits and demography to explain complex temperature responses across levels of organization,” Proceedings of the National Academy of Sciences, vol. 118, no. 42, p. e2104863118, 2021.

[17] M. J. Follows, S. Dutkiewicz, S. Grant, and S. W. Chisholm, “Emergent biogeography of microbial communities in a model ocean,” science, vol. 315, no. 5820, pp. 1843–1846, 2007.

[18] B. A. Ward, S. Dutkiewicz, O. Jahn, and M. J. Follows, “A size-structured food-web model for the global ocean,” Limnology and Oceanography, vol. 57, no. 6, pp. 1877–1891, 2012.

[19] E. Litchman and C. A. Klausmeier, “Trait-based approaches to phytoplankton ecology,” Annual Review of Ecology, Evolution, and Systematics, vol. 39, pp. 615–639, 2008.

[20] G. B. West, J. H. Brown, and B. J. Enquist, “A general model for the origin of allometric scaling laws in biology,” Science, vol. 276, no. 5309, pp. 122–126, 1997.

[21] J. H. Brown, J. F. Gillooly, A. P. Allen, V. M. Savage, and G. B. West, “Toward a Metabolic Theory of Ecology,” Ecology, vol. 85, no. 7, pp. 1771–1789, 2004. eprint: https://onlinelibrary.wiley.com/doi/pdf/10.1890/03-9000.

[22] I. A. Hatton, A. P. Dobson, D. Storch, E. D. Galbraith, and M. Loreau, “Linking scaling laws across eukaryotes,” Proceedings of the National Academy of Sciences, vol. 116, pp. 21616–21622, Oct. 2019.

[23] H. Hillebrand, E. Acevedo-Trejos, S. D. Moorthi, A. Ryabov, M. Striebel, P. K. Thomas, and M.-L. Schneider, “Cell size as driver and sentinel of phytoplankton community structure and functioning,” Functional Ecology, vol. 36, no. 2, pp. 276–293, 2022.

[24] E. Marañón, “Cell Size as a Key Determinant of Phytoplankton Metabolism and Community Structure,” Annual Review of Marine Science, vol. 7, pp. 241–264, Jan. 2015.

[25] I. Gallego, P. Venail, and B. W. Ibelings, “Size differences predict niche and relative fitness differences between phytoplankton species but not their coexistence,” The ISME Journal, vol. 13, pp. 1133–1143, 01 2019.

[26] A. J. Lotka, Elements of Physical Biology. Williams & Wilkins, 1925.

[27] V. Volterra, Variazioni e fluttuazioni del numero d’individui in specie animali conviventi. Societá anonima tipografica” Leonardo da Vinci”, 1926.

[28] L. Fant and G. Ghedini, “Biomass competition connects individual and community scaling patterns,” Nature Communications, vol. 15, p. 9916, Nov. 2024.

[29] M. Barbier, C. de Mazancourt, M. Loreau, and G. Bunin, “Fingerprints of high-dimensional coexistence in complex ecosystems,” Phys. Rev. X, vol. 11, p. 011009, Jan 2021.

[30] Y. R. Zelnik, N. Galiana, M. Barbier, M. Loreau, E. Galbraith, and J.-f. Arnoldi, “How collectively integrated are ecological communities?,” Ecology Letters, vol. 27, no. 1, p. e14358, 2024.

[31] J. Wickman, E. Litchman, and C. A. Klausmeier, “Eco-evolutionary emergence of macroecological scaling in plankton communities,” Science, vol. 383, no. 6684, pp. 777–782, 2024.

[32] S. M. Ernest, B. J. Enquist, J. H. Brown, E. L. Charnov, J. F. Gillooly, V. M. Savage, E. P. White, F. A. Smith, E. A. Hadly, J. P. Haskell, et al., “Thermodynamic and metabolic effects on the scaling of production and population energy use,” Ecology Letters, vol. 6, no. 11, pp. 990–995, 2003.

[33] J. DeLong and D. Hanson, “Metabolic rate links density to demography in,” 2009.

[34] L. Schuster, H. Cameron, C. R. White, and D. J. Marshall, “Metabolism drives demography in an experimental field test,” Proceedings of the National Academy of Sciences, vol. 118, no. 34, p. e2104942118, 2021.

[35] E. Marañón, P. Cermeño, D.C. López-Sandoval, T. Rodríguez-Ramos, C. Sobrino, M. Huete-Ortega, J. M. Blanco, and J. Rodríguez, “Unimodal size scaling of phytoplankton growth and the size dependence of nutrient uptake and use,” Ecology Letters, vol. 16, no. 3, pp. 371–379, 2013.

[36] M. Czarnoleski and W. C. Verberk, “Cell size matters: a unifying theory across the tree of life,” Trends in Ecology Evolution, vol. 40, no. 11, pp. 1113–1125, 2025.

[37] J. Hinners, P. A. Argyle, N. G. Walworth, M. A. Doblin, N. M. Levine, and S. Collins, “Multi-trait diversification in marine diatoms in constant and warmed environments,” Proceedings of the Royal Society B: Biological Sciences, vol. 291, no. 2019, 2024.

[38] J. E. Goldford, N. Lu, D. Bajić, S. Estrela, M. Tikhonov, A. Sanchez-Gorostiaga, D. Segré, P. Mehta, and A. Sanchez, “Emergent simplicity in microbial community assembly,” Science, vol. 361, no. 6401, pp. 469–474, 2018.

[39] G. Ghedini, D. Marshall, and M. Loreau, “Phytoplankton diversity affects biomass and energy production differently during community development,” Functional Ecology, vol. 36, 2022.

[40] H. Hillebrand, C.-D. Dürselen, D. Kirschtel, U. Pollingher, and T. Zohary, “Biovolume calculation for pelagic and benthic microalgae,” Journal of phycology, vol. 35, no. 2, pp. 403–424, 1999.

[41] S. Arya, A. B. George, and J. O’Dwyer, “The architecture of theory and data in microbiome design: towards an s-matrix for microbiomes,” Current Opinion in Microbiology, vol. 83, p. 102580, 2025.

[42] R. R. Guillard and J. H. Ryther, “Studies of marine planktonic diatoms: I. cyclotella nana hustedt, and detonula confervacea (cleve) gran.,” Canadian journal of microbiology, vol. 8, no. 2, pp. 229–239, 1962.

[43] H. Hillebrand, C.-D. Dürselen, D. Kirschtel, U. Pollingher, and T. Zohary, “Biovolume calculation for pelagic and benthic microalgae,” Journal of phycology, vol. 35, no. 2, pp. 403–424, 1999.

