## supplementary materia for "Ecological predictability emerges at the population level in phytoplankton communities"

### Supplementary Figures and Tables

**Table S1. Species and reference strains used in the experiments.**

| Experiment Location | Culture collection | Species name | Strain code | Group |
|---|---|---|---|---|
| Australia | ANACC (Austalia) | *Amphidinium carterae* | CS-740 | Dinoflagellata |
| | | *Tetraselmis* sp. | CS-91 | Chlorophyta |
| | | *Dunaliella tertiolecta* | CS-14 | Chlorophyta |
| | | *Tisochrysis lutea* | CS-177 | Haptophyta |
| | | *Synechococcus* sp. | CS-94 | Cyanobacteria |
| Portugal | Roscoff (France) | *Amphidinium carterae* | RCC88 | Dinoflagellata |
| | | *Dunaliella tertiolecta* | RCC6 | Chlorophyta |
| | | *Phaeodactylum tricornutum* | RCC2967 | Bacillariophyta |
| | | *Tisochrysis lutea* | RCC90 | Haptophyta |
| | | *Nannochloropsis granulata* | RCC438 | Eustigmatophyceae |

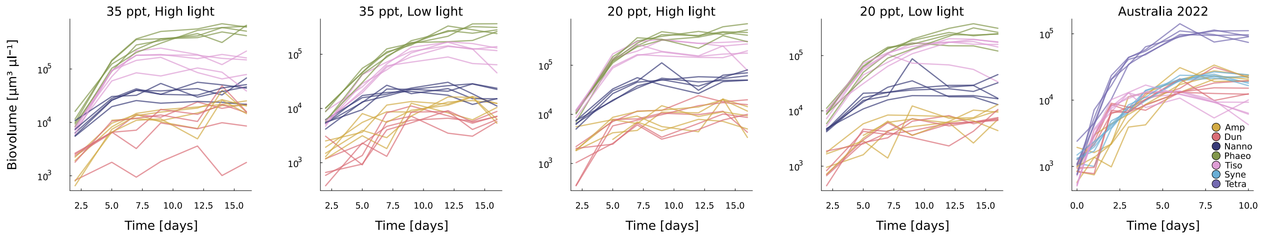


**Fig. S1.** Community dynamics is reproducible across samples with the same environmental conditions. Growth trajectories of each species in phytoplankton communities, within each environmental condition.

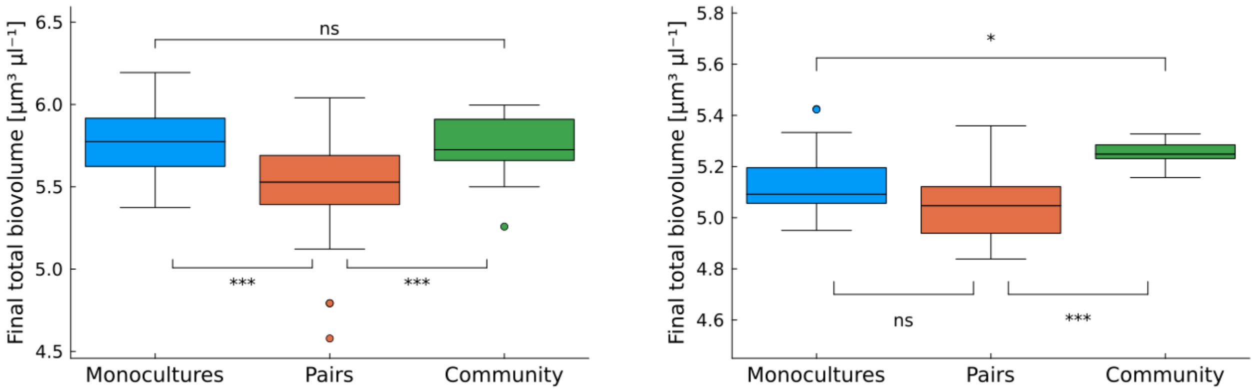


**Fig. S2.** Pairs of species grow to lower densities than monocultures and communities of five species. Comparison of final total biovolume across monocultures, pair cultures, and multispecies communities at the two experimental locations. Pair cultures consistently reach lower total biomass than monocultures, motivating the rescaling used when estimating pairwise interaction coefficients.

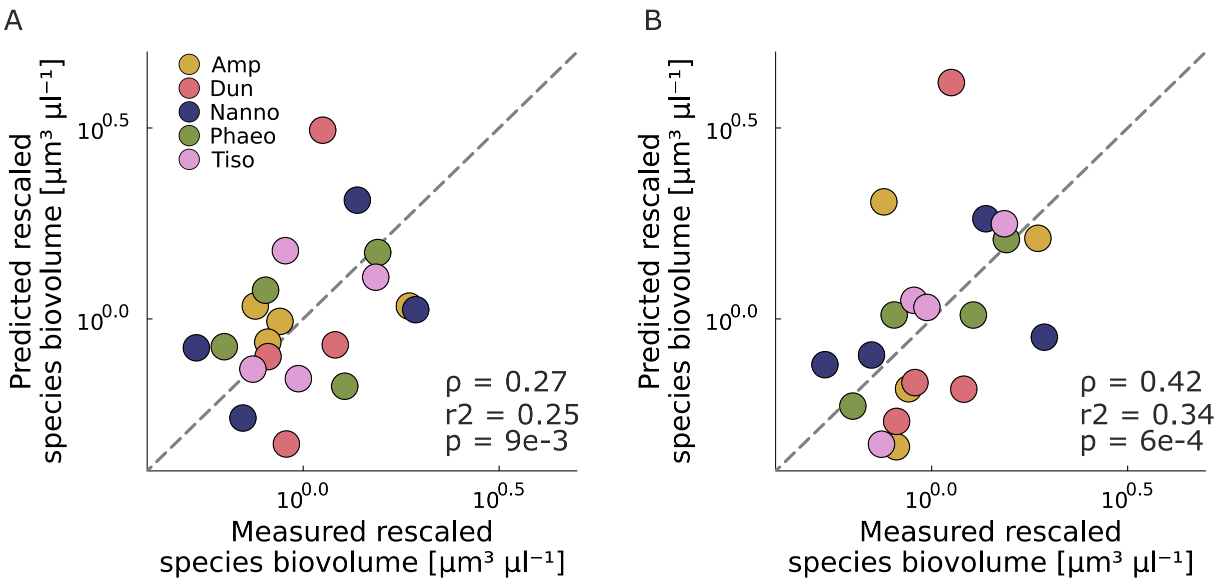


**Fig. S3.** Interactions improve the prediction accuracy across treatments in PT data. Comparison of final biovolume predicted and measured in monocultures. Data are rescaled by the average species biomass density across environmental conditions. The introduction of species interactions (B) increases the correlation between measures and predictions across environmental conditions for all species

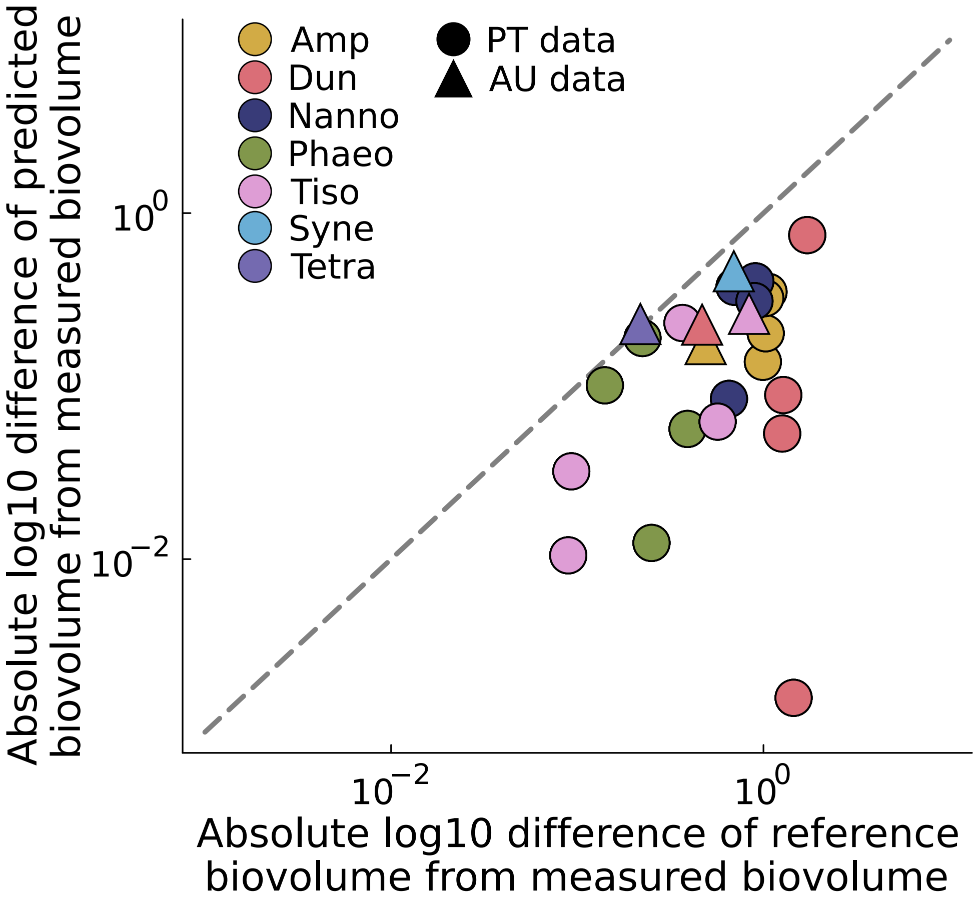


**Fig. S4.** Interactions with more than one species are necessary to explain community composition. Comparison of the difference between the $log_{10}$ average final biovolume predicted and measured in monocultures and the difference between the $log_{10}$ average final biovolume of pairs (i.e. reference biovolume, rescaled by the factor $c$) and the average measured biovolume

### References
